# Variations in human trigeminal and facial nerve branches and foramina identified by dissection and microcomputed tomography

**DOI:** 10.1101/2024.09.17.613414

**Authors:** Hannah L. Grimes, Valentina Pizzuti, Maria A. Wright, Thomas Santarius, Susan Jones

## Abstract

The aim of this study was to identify branches of the trigeminal and facial nerves relevant to surgical incisions and injections and the scalp block techniques in the frontotemporal region, and to determine their relationships with superficial vascular structures and bony landmarks. Half-heads from consenting embalmed donors (6 male, 2 female, mean age at death 78.4 years) were used in this study. Detailed dissection was carried out to identify the position of the auriculotemporal nerve (ATN) relative to the superior temporal artery (STA) and the facial nerve (FN) in six subjects (5 male, 1 female). The results provide a minimum safe distance of 5 mm between the STA and the frontotemporal branches of the FN at the level of the low edge of zygoma and 8mm between the low edge of zygoma and the FN trunk, providing a pre-auricular triangle of safety for incisions and injections. Variability between subjects was up to 60%. Microcomputed tomography (microCT) scans were taken from all eight subjects and the three-dimensional reconstructions were used to identify the supraorbital notch (SON), the zygomaticotemporal foramen (ZTF) and the zygomaticofacial foramen (ZFF). The volume and relative locations of these foramina were calculated for 5-8 subjects. The closest distance between ZTF and the FZS ranged from 9 to 21mm (26% variation); 3 subjects had a single ZTF while 5 subjects had two ZTF. The angle at the centre of the orbit between ZFF and the FZS ranged from 156 to 166 degrees (2.5% variation). These findings demonstrate that both traditional cadaveric dissection methods as well as contemporary microCT methods can be used to investigate the relative locations of nerves or their foramina in the human head. The findings provide anatomical considerations for fronto-temporal incisions and local anaesthesia.

## Introduction

Knowledge of the location and course of nerves in the frontotemporal region is key to safe and effective surgery carried out by neurosurgical, maxillofacial, ophthalmic and plastic surgeons. Regional scalp block is a technique that involves injection of local anaesthetic at well-defined anatomical points, targeting the major sensory innervation of the face and scalp [1]. It is used to facilitate intra-operative brain mapping while permitting communication with patients during surgical procedures for epilepsy, tumour resection and vascular conditions [1-6]. Local anaesthetic infiltration of the scalp before craniotomy is effective in reducing tachycardia and hypertension, which could otherwise result in increased cerebral blood flow and intracranial pressure [7-10].

Moreover, local anaesthetic infiltration may also prevent the need for increased analgesic requirements during the surgical procedure and reduce postoperative pain and opioid analgesic requirements [8, 10-13].

Sensory innervation of the scalp is provided by both the trigeminal and spinal nerves. The trigeminal nerve has ophthalmic, maxillary and mandibular divisions, all of which contribute branches that innervate the forehead and scalp [14-16]. The nerves targeted in a scalp block are the supraorbital and supratrochlear branches of the ophthalmic division, the auriculotemporal and zygomaticotemporal branches of the maxillary division, as well as the greater and lesser occipital nerves. Block of the zygomaticotemporal nerve, as well as the zygomaticofacial nerve is required for some maxillofacial procedures [17-19]. During regional scalp block there are currently two main potential complications for patients. Firstly, insufficient block of the relevant nerve branches may lead to pain and, consequently, substandard or unsafe awake surgery. Secondly, scalp block can result in transient, facial nerve palsy [20-22], which can be worrying for both the patient and the surgeon, especially if one is not sufficiently familiar with the anatomy. Identification of optimal injection sites is important to ensure maximum benefit whilst avoiding neural structures which may carry adverse post-operative outcomes. The aim of this study was to test two different approaches to locating trigeminal nerve branches: traditional cadaveric dissection of the ATN and FN and microCT to locate SON, ZTF and ZFF. These approaches were used to identify the relationships between the trigeminal nerve branches or their foramina with palpable structures.

Evidence is presented that a combined approach can provide insight into the location of trigeminal nerve branches and that there is anatomical variation.

## Methods

Adult half-heads (right side) from 8 consenting donors (6 male, 2 female; mean age 78.4 years at death) were used in this study. Donors were embalmed using a solution composed of ethanol (38%), formaldehyde (5%) and distilled water. Demographic and clinical data collected for each donor are shown in Table 1. Detailed dissection was performed with 6 subjects (5 male, 1 female) to isolate the auriculotemporal branch of the trigeminal nerve. Skin, subcutaneous fat and connective tissue were carefully removed, retaining the relationship of the ATN to other structures, including branches from the facial nerve trunk (FNT), zygoma and STA. Dissections were performed in the Human Anatomy Centre of the Department of Physiology, Development & Neuroscience, University of Cambridge, and then photographed. Measurements of distances between major branches of trigeminal and facial nerves and facial landmarks were made on the specimens by two independent investigators using micro-calipers and the average values are reported (Table 2).

**Table 1.** Demographic and clinical data for all donors.

| Subject | Gender | Age (yr) | Cause of death |
| --- | --- | --- | --- |
| 1 | M | 67 | Glioblastoma |
| 2 | M | 97 | Metastatic carcinoma |
| 3 | M | 76 | COPD |
| 4 | F | 71 | Astrocytoma |
| 5 | M | 90 | Stroke/ pneumonia |
| 6 | M | 78 | COPD |
| 7 | F | 57 | Unknown primary/metastasis to lungs |
| 8 | M | 91 | Lung and kidney cancer |

**Table 2.** Shortest distances to the indicated nerve branches (mm). STA_Z_ and dFN_Z_ are the superior temporal artery and the most dorsal branch of the facial nerve respectively, both measured at lower edge of the zygoma. ATN is the auriculotemporal nerve, FNT is the facial nerve trunk, and pdFN is the most posterior part of the dorsal facial nerve branch. ND, not determined.

| **Subject** | **STA<sub>Z</sub>-dFN<sub>Z</sub>\***<br>***AP*** | **STA<sub>Z</sub>\* –**<br>**pdFN *CC*** | **STA-ATN**<br>***AP*** | **ATN – FNT**<br>***CC*** | **Tragus apex -**<br>**FNT *CC*** |
| --- | --- | --- | --- | --- | --- |
| 1 | 13 | 14 | 3 | 28 | 14 |
| 2 | 5 | 8 | 4 | *ND* | 18 |
| 3 | 6 | 17 | 4 | 21 | 21 |
| 4 | 16 | 19 | 4 | 22 | 21 |
| 5 | 17 | 12 | 5 | 17 | 19 |
| 6 | 27 | 24 | 4 | 27 | 29 |
| **Mean** | **14** | **16** | **4** | **23** | **20** |
| **SD** | **8** | **6** | **1** | **4** | **5** |
| **CV** | **0.6** | **0.4** | **0.2** | **0.2** | **0.2** |

MicroCT scans were taken from all eight subjects using a Nikon XT H 225 cone-beam scanner and 1080 projection angles, with two frames averaged per projection and an exposure time of 1000 milliseconds. Each scan took approximately 40 minutes. This was to balance the resolution of the tomograms against the risk of specimens shifting during the scan. Settings of 170-220 μA and 175- 200 kV were used, and a 0.5 mm copper filter was used in most cases. Voxel sizes ranged from 0.0624-0.0744 mm. Around 1800 tomograms per scan were produced as TIF files. One third of these were used to generate a three-dimensional reconstruction using MicroView (version 2.5) software, for visualising the surface and sectional locations of the supraorbital notch (SON), the zygomaticotemporal foramen (ZTF) and the zygomaticofacial foramen (ZFF). TIF files were converted to JPG files using GIMP software (version 2.10) for importing into Stradwin (version 6.0) or StradView (Version 6.02) which allowed sectioning of channels within the bone. From this, estimates of volume for each channel and their relative positions were obtained (defined in Table 3). Sectioning consistency was verified by two independent investigators for the SON (5 subjects; mean difference 10 ± 6.3%) and the ZFF (3 subjects; mean difference 1.7 ± 3.2%).

**Table 3.** Boundaries of each bone channel measured using 3D reconstruction from microCT images. Only direct channels from internal to external boundaries were sectioned. SON, supra- orbital notch; ZFF, zygomaticofacial foramen; ZTF, zygomaticotemporal foramen.

| Subject | SON | ZFF | ZTF |
| --- | --- | --- | --- |
| Definition | Channel through the bone for the supra-orbital and supra-trochlear nerves | Channel through the bone for the zygomaticofacial nerve | Channel through the bone for the zygomaticotemporal nerve |
| External boundary | Superior to the orbit; nasal side | Opening onto the anterior zygoma, lateral to or below the orbit | Opening onto the posterior zygoma, in the lateral margin of the orbit |
| Positional information | Angle formed at centre of orbit between SON and FZS | Angle formed at centre of orbit between ZFF and FZS | Shortest distance between ZTF and FZS |

## Results

The ATN and FNT were identified and dissected in 6 cadaveric specimens from five male donors and one female donor. In order to determine a pre-auricular triangle of safety, similar to the preauricular safe zone described by Kucukguven et al (2021) [24], for incisions and injections, where no nerves or vessels are likely to be present, two measurements were made (Figure 1; Table 2). The anteroposterior distance from the STA to the frontotemporal branch of the FN (where both cross the lower edge of zygoma: STA_Z_ and ftFN_Z_) ranged from 5 to 27mm, with 60% variation between subjects. The craniocaudal distance from the STA_Z_ to the most posterior part of the FNT (pFN) ranged from 8 to 24mm with 40% variation between subjects. The shortest following distances were also measured for each subject (Figure 2; Table 2). The anteroposterior distance from ATN to the STA ranged from 3-5 mm; ATN was posterior to STA in all 6 subjects. The craniocaudal distance from ATN to FNT ranged from 17 to 28 mm). Tragus apex to the FNT in the craniocaudal axis ranged from 14 to 29 mm. The extent of variability between subjects was 20% for these three measurements.

**Figure 1:**
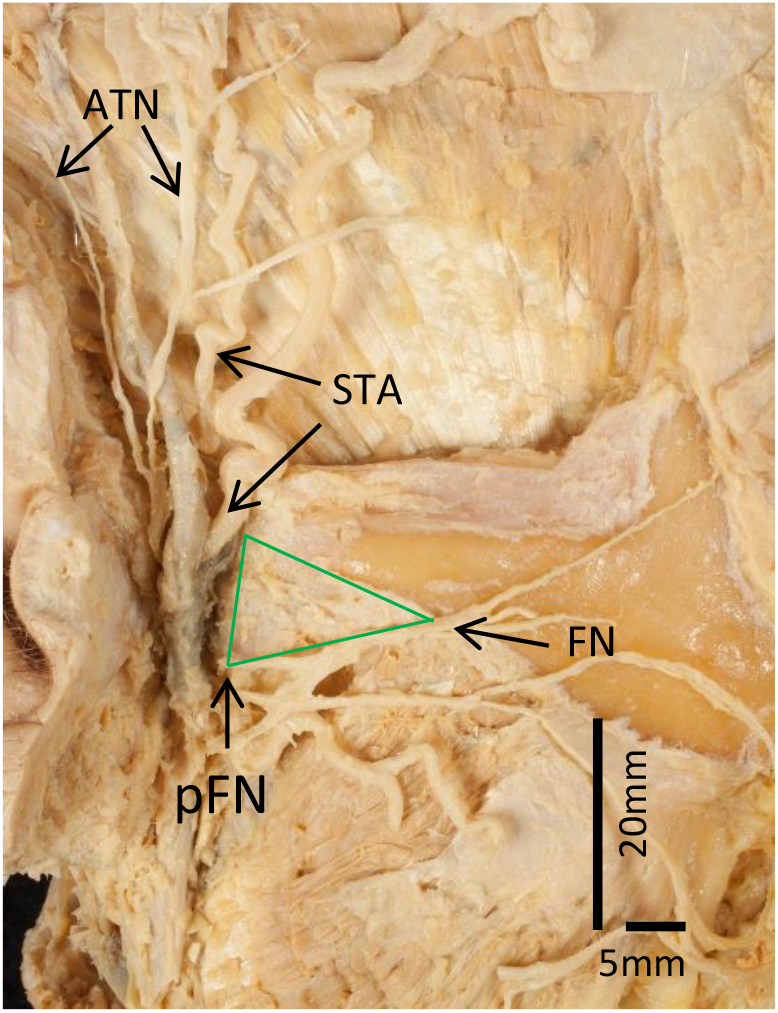
Example dissection of auriculotemporal nerve (ATN) and facial nerve (FN) branches in subject 5 (male) to show a pre-auricular triangle of safety (green). A, distance from STA to frontotemporal branch of FN at lower edge of zygoma (Z) (17mm); B, distance from STA at lower edge of zygoma to most posterior part of FN (pFN) (12mm). STA/ vein: superficial temporal artery/ vein.

**Figure 2:**
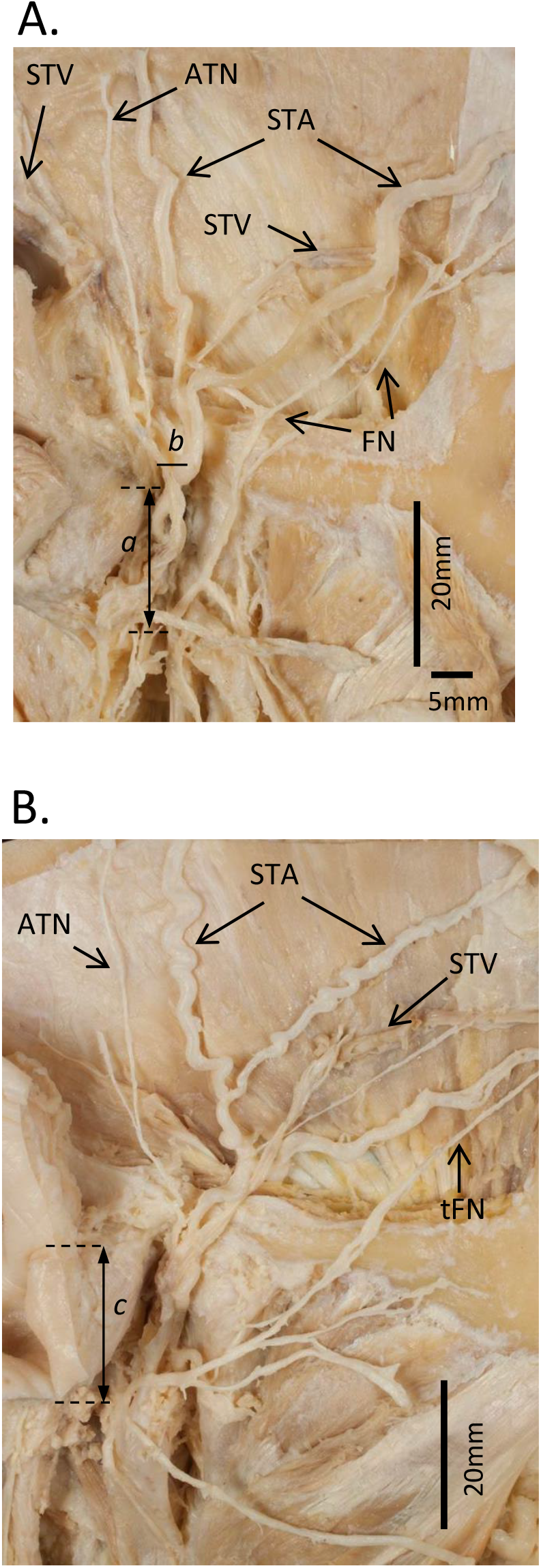
A. Example dissection of auriculotemporal nerve (ATN) and frontotemporal facial nerve (FN) branches in subject 3 (male): *a* indicates the distance from the ATN to FNT (21mm); *b* indicates the distance from the ATN to the STA (4mm). B. Example dissection in subject 4 (female): *c* indicates the distance from tragus apex to the facial trunk (21mm). STA/V: superficial temporal artery/ vein.

Using tomograms obtained from microCT scanning, reconstructions of the skull bones housing the SON, the ZTF and the ZFF were made in MicroView from one third of the tomograms (examples shown in Figure 3A, B; all subjects shown in Figure 4) and in Stradwin from the full set of tomograms (examples shown in Figure 3C, D). The sections through the SON (defined in Table 3; indicated in Figure 3C) were examined in 8 subjects and a brief description is given in Table 4. The volume of the SON was determined in 7 subjects by sectioning using Stradwin software and ranged from 0.008 to 0.056 ml^3^ (CoV: 0.85; Table 4); the female subjects had the smallest (subject 7) and the largest (subject 4) SON volumes. Using Stradwin, the angle formed between the (clinically palpable) SON and FZS at the centre of the orbit was determined for six subjects (Table 4; method illustrated in Figure 5A) and ranged from 80 to 99 degrees (mean ± SD: 91 ± 7; CoV: 0.08; range of angles illustrated in Figure 5; female subject 4 was in the middle of the range). Thus, while the volume measurements varied considerably, the angle measurement was much less variable between subjects, and there was no obvious relationship between sex and these measurements.

**Table 4.** Description, volume measurement and positional measurement for the supraorbital notch (SON) in each subject. Mean, standard deviation and coefficient of variation shown in bold. ND, not determined.

| **Subject** | **SON description** | **SON volume**<br>**(ml<sup>3</sup>)** | **Angle of SON to FZS**<br>**(degrees)** |
| --- | --- | --- | --- |
| 1 | True notch | 0.018 | 80 |
| 2 | Almost fully open<br>foramen | 0.021 | *ND* |
| 3 | Almost fully open<br>foramen | 0.012 | 99 |
| 4 | Almost fully open<br>foramen | 0.056 | 92 |
| 5 | Partially open notch | *ND* | 86 |
| 6 | Fully open foramen | 0.08 | 98 |
| 7 | True notch | 0.008 | *ND* |
| 8 | Fully open foramen | 0.023 | 88 |
|  | **Mean** | **0.031** | **90.5** |
|  | **SD** | **0.027** | **7.3** |
|  | **CoV** | **0.85** | **0.08** |

**Figure 3:**
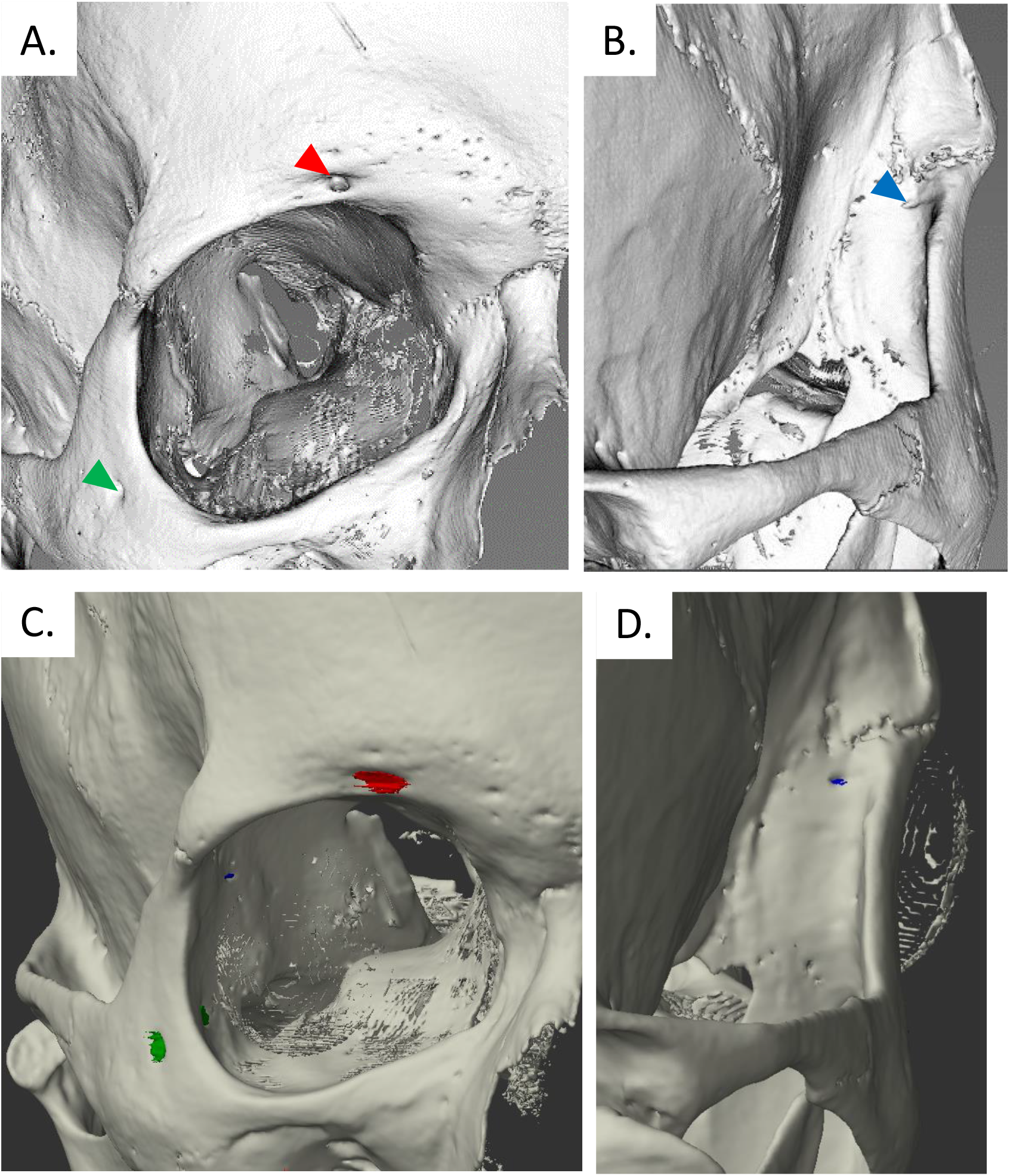
A. Reconstruction from one third of all micro computed tomograms collected from subject 1 (male), generated in MicroView. Red arrow indicates the supraorbital notch, green arrow the zygomaticofacial foramen. B. MicroView reconstruction showing the zygomaticotemporal foramen (blue arrow). C. Reconstruction of the sectioned foramina and the facial bones generated in Stradwin showing the supraorbital notch (red) and the zygomaticofacial foramen (green). D. Stradwin reconstruction showing the zygomaticotemporal foramen (blue).

**Figure 4:**
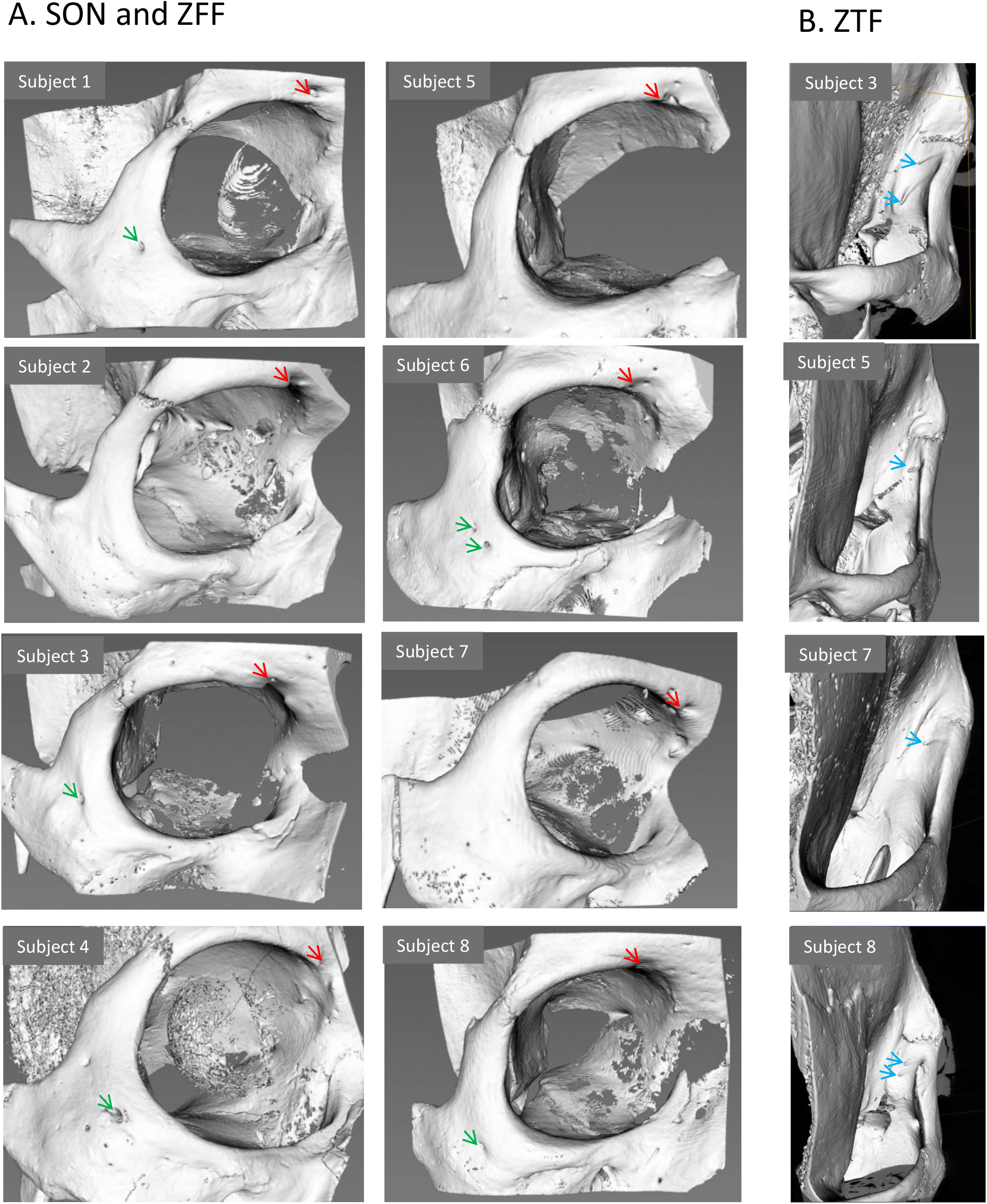
Full reconstruction from one third of all tomograms generated in MicroView. A. Supraorbital notch and zygomaticofacial foramen from all subjects. B. Zygomaticotemporal foramen from 4 subjects.

**Figure 5:**
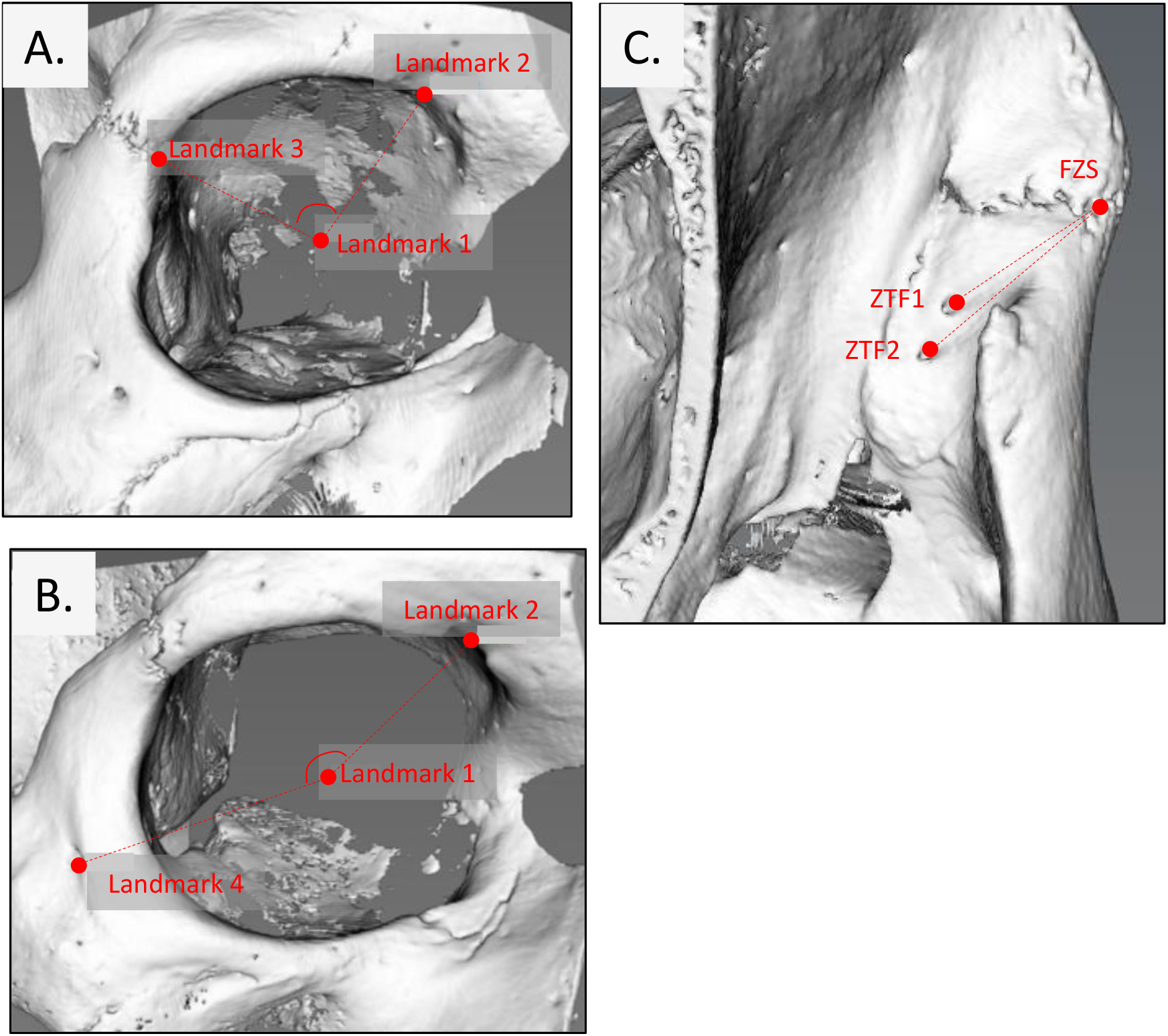
Illustration of the landmark method used to determine angles and distances. A. The angle formed by the supraorbital notch (SON, landmark 2) and the frontozygomatic suture (FZS, landmark 3) at the centre of the orbit (CoO, landmark 1) in subject 6 (male). B. The angle formed by the zygomaticofacial foramen (landmark 4) and the SON (landmark 2) at the CoO (landmark 1) in subject 3 (male). C. The shortest distance of the zygomaticotemporal foramen (landmark 5) to the outer edge of the FZS (landmark 3) in subject 8 (male).

MicroCT was also used to assess the position of the foramina of the zygomaticotemporal nerve, targeted in scalp block, and zygomaticofacial nerve, also relevant to maxillofacial surgery. The ZTF (defined in Table 3; examples shown in Figure 3D and Figure 4) was examined in 8 subjects and a brief description is given in Table 5; 3 subjects had a single exterior opening and 5 subjects had two exterior openings. ZTF volume measurements from 6 subjects (including both channels when present) ranged from 0.002 to 0.019 ml^3^ (mean ± SD: 0.005 ± 0.003 ml^3^; CoV: 0.65; Table 5). Both female subjects were in the middle of this range. The distance between the ZTF and the FZS was determined for 6 subjects (Table 5; method illustrated in Figure 5C); when accounting for only the closest of two ZTF (where present), distances ranged from 9 to 21mm (15 ± 4m; CoV: 0.26). Thus, the size, number of exterior openings and location of the ZTF relative to FZS varied between subjects, and there was no obvious relationship between sex and the number, volume or position of the ZTF.

**Table 5.** Description, volume measurement and positional measurement for the zygomaticotemporal foramen (ZTF) in each subject. Mean, standard deviation and coefficient of variation shown in bold. ND, not determined.

| **Subject** | **ZTF description** | **ZTF volume**<br>**(ml<sup>3</sup>)** | **Distance of**<br>**closest ZTF to FZS**<br>**(mm)** |
| --- | --- | --- | --- |
| 1 | One exterior opening | 0.002 | 14 |
| 2 | Two exterior openings | 0.008 | 17 |
| 3 | Two exterior openings | 0.006 | 15 |
| 4 | Two exterior openings | 0.009 | 21 |
| 5 | One exterior opening | *ND* | *ND* |
| 6 | Two exterior openings | 0.019 | 14 |
| 7 | One exterior opening | 0.004 | 9 |
| 8 | Two exterior openings | *ND* | *ND* |
|  | **Mean** | **0.005** | **15** |
|  | **SD** | **0.003** | **3.9** |
|  | **CoV** | **0.65** | **0.26** |

The ZFF (defined in Table 3; example shown in Figure 3C, all subjects shown in Figure 4) was examined in 8 subjects and a brief description is given in Table 6. In 3 subjects there was no obvious exterior opening or channel from exterior to interior in the expected location of ZFF, therefore volume measurements were made from 5 subjects and ranged from 0.011 (in female subject 4) to 0.046 ml^3^ (mean ± SD: 0.031± 0.013 ml^3^; CoV: 0.4; Table 6). The angle formed between the ZFF and the SON at the centre of the orbit was determined for 5 subjects (Table 6; method illustrated in Figure 5B) and ranged from 156 to 166 (in female subject 4) degrees (mean ± SD: 159 ± 4; CoV: 0.025; range of angles illustrated in Figure 5). As for SON, there was considerable variation in the volume measurements while the angle measurement was much less variable between subjects.

**Table 6.** Description, volume measurement and positional measurement for the zygomaticofacial foramen (ZFF) in each subject, where present. Mean, standard deviation and coefficient of variation shown in bold. NA, not available.

| **Subject** | **ZFF description** | **ZFF volume**<br>**(ml<sup>3</sup>)** | **Angle of ZFF to**<br>**SON**<br>**(degrees)** |
| --- | --- | --- | --- |
| 1 | One exterior opening | 0.046 | 157 |
| 2 | *No obvious exterior opening or*<br>*channel to interior: not*<br>*sectioned* | NA | NA |
| 3 | One exterior opening | 0.035 | 156 |
| 4 | One exterior opening | 0.011 | 166 |
| 5 | *No obvious exterior opening or*<br>*channel to interior: not*<br>*sectioned* | NA | NA |
| 6 | Two exterior openings | 0.03 | 160 |
| 7 | *No obvious exterior opening or*<br>*channel to interior: not*<br>*sectioned* | NA | NA |
| 8 | One exterior opening | 0.032 | 158 |
|  | **Mean** | **0.031** | **159.4** |
|  | **SD** | **0.013** | **4.0** |
|  | **CoV** | **0.4** | **0.025** |

## Discussion

Detailed dissection and microCT were used to examine the relationships between neural, vascular and bony structures of the face in order to assess the variations in trigeminal nerve branches. Our findings demonstrate that a combination of cadaveric dissection of large nerve branches combined with microCT for the foramina of finer nerve branches can be used to assess the variability in the likely locations of trigeminal nerve branches. The data show that there is anatomical variation in trigeminal nerve branches that are targeted in regional scalp block and this needs to be taken into account when planning surgical approaches. Understanding these variations will help minimise adverse consequences for patients. We propose a pre-auricular triangle of safety, similar to the preauricular safe zone described by Kucukguven et al (2021) [24], for making injections during scalp block that are less likely to damage nerves or vessels.

The ATN innervates the skin of the temple. In this study there was 20% variation in the distance between the ATN and the STA, although the ATN was posterior to STA in all 6 subjects. Knowing the range distances of the ATN from STA will be helpful when making incisions for temporal and pterional craniotomies. To avoid injury to the main trunk of STA, the incision is best placed just anterior to the STA. However, as only 1 out of 6 subjects had an ATN-STA distance of <4 mm; based on this, when placing the incision within 4mm posterior to STA the chances of damaging the ATN would be less than 17%. This may be a relevant consideration, if the posterior branch of the STA is to be preserved, such as during cerebral flow augmentation procedures [25] The spatial association of the ATN with the temporal, zygomatic, and buccal branches of the facial nerve (at the border of the masseter) places all these branches at risk. For example, transient facial paralysis lasting several hours can sometimes be seen following ATN nerve block [20-23] which can be explained by close proximity of the ATN and the facial nerve. In this study, the shortest distance between the ATN and the FNT was 17mm and so variations here should cause no real problems during regional nerve block. At the level of the lower edge of zygoma, the minimum distance between the STA and the frontotemporal branches of the trigeminal nerve was 5mm and the shortest distance from STA to the posterior facial nerve was 8mm, therefore the pre-auricular triangle of safety is a small area in some subjects. In a study of 20 Caucasian subjects the pre-auricular safe zone was 10mm [24], a little broader than our minimum distance of 5mm. The combined output of these studies highlights and refines our understanding of the variability of the relevant neurovascular structures in this area, making anaesthesia and surgery safer.

The supraorbital and supratrochlear nerves are branches of the ophthalmic division of the trigeminal nerve and they pass via the SON to supply the skin over the upper eyelid and the forehead, respectively. A landmark method was used to quantify the location of the SON relative to the FZS by measuring the angle formed between these structures at the orbital rim with the apex at the centre of the orbit. Interestingly, less than 10% variation in this angle was observed between subjects. A similar approach was used to quantify the location of the ZFF; the angle formed at the centre of the orbit between ZFF and SON showed less than 3% variation between subjects.

Triangulation methods could potentially be incorporated into the anaesthetic technique for consistently and accurately finding the position of targeted nerves. ZFF was absent in three of the eight subjects. This is rare, but it has been described previously [26]. Although nerve block of ZFF is not usually required for awake craniotomies, it is a useful landmark for skull base, maxillofacial and oculoplastic surgeries.

The zygomaticotemporal nerve supplies the skin over the cheek. In this study, there was variation in the number of exterior openings of the ZTF, and 26% variation in the distance between the FZS and the closest opening. Loukas et al [26] studied the location of the ZTF on human dry skulls (121 male and 79 female); in half of the specimens, the zygomaticotemporal foramen was not present in the zygomatic bone. Another study on 192 Korean skulls noted an absence of the zygomaticotemporal foramen in only 7.3% of skulls examined [27]. This dramatically different result was considered a result of size differences between zygomatic bones from those of Asian compared with European descent, although some of this discrepancy can probably be attributed to the definition of the ZTF by Loukas et al. This reinforces the argument that determination of the optimum location for block sites requires the anatomical variation seen between different ethnic groups to be examined in more depth, and that diversity should be a focus in future research [28]. Although damage to this nerve is inevitable on most frontolateral and pterional approaches, the nerve can be spared in some, especially temporal, craniotomies if the surgeon is aware of the location of the nerve foramina.

## Conclusions

Detailed knowledge of the variations of the anatomical locations of the nerves is important for safe local anaesthesia and surgery in the frontotemporal region. We described a pre-auricular triangle of safety, which may be helpful in planning anaesthetic injections and surgical incisions. To overcome the limitations of conventional anatomical dissections we employed novel segmentation technique and microcomputed tomography to accurately describe the anatomical locations of zygomaticotemoporal and zygomaticofacial nerves and their variations. We have shown that when using the landmark method of angles and distances, the relative locations of ZFS, ZON and ZFF show surprisingly little variance, which can be of clinical use. Small cohorts and data sets limit the conclusions we can draw but we have shared our methods and findings in order that further work can help to progress towards more powerful conclusions.

## Abbreviations

ATN: auriculotemporal nerve
STA: superior temporal artery
FN(T): Facial nerve (trunk)
SON: supraorbital notch
ZTF: zygomaticotemporal foramen
ZFF: zygomaticofacial foramen
FZS: frontozygomatic suture
*AP*: anterior to posterior
*CC*: cranial to caudal
microCT: Microcomputed tomography

## Acknowledgments

The authors thank the Cambridge Biotomography Centre for the use of their scanner, Ket Smithson for technical support and Professor Matthew Mason for invaluable help with scanning and essential advice on analysis of microCT images.

